# Inkjet-printed transparent electrodes for electrical brain stimulation

**DOI:** 10.1101/2024.09.06.611618

**Authors:** Rita Matta, Davide Reato, Alberto Lombardini, David Moreau, Rodney P. O’Connor

**Author notes:** Equal contribution (Rita Matta, Davide Reato). Equal contribution (David Moreau, Rodney P. O’Connor).

## Abstract

Electrical stimulation is a powerful tool for investigating and modulating brain activity, as well as for treating neurological disorders. However, understanding the precise effects of electrical stimulation on neural activity has been hindered by limitations in recording neuronal responses near the stimulating electrode, such as stimulation artifacts in electrophysiology or obstruction of the field of view in imaging. In this study, we introduce a novel stimulation device fabricated from conductive polymers that is transparent and therefore compatible with optical imaging techniques. The device is manufactured using a combination of microfabrication and inkjet printing techniques and is flexible, allowing better adherence to the brain’s natural curvature. We characterized the electrical and optical properties of the electrode and evaluated its performance in the brain of an anesthetized mouse. Furthermore, we combined experimental data with a finite-element model of the in-vivo experimental setup to estimate the maximum electric field that the highly transparent device can generate in the mouse brain. Our findings indicate that the device can generate an electric field as high as 300 V/m, demonstrating its potential for studying and manipulating neural activity using a range of electrical stimulation techniques relevant to human applications. Overall, this work presents a promising approach for developing versatile new tools to apply and study electrical brain stimulation.

## INTRODUCTION

Electrical brain stimulation is a long-established technique for studying brain function [1,2]. In addition to its research applications, electrical stimulation has been used clinically to treat or alleviate symptoms of various neurological conditions, either using intracranial or transcranial techniques [3–6], as well as for neuroprosthetic purposes [7,8].

Thanks to insights from computational models and in-vitro and in-vivo experiments, we now have a relatively precise understanding of the electric field distributions in the brain generated by different electrical stimulation techniques [9,10], as well as their effects on individual neurons [11–13]. The primary direct effect of electrical stimulation on neurons is a change in membrane potential [14,15], which depends on several factors, including the direction and magnitude of the electric field, the specific waveform used, and the morphology and electrophysiological properties of the cell [16–19]. If the electric field is sufficiently strong (>20 V/m), membrane voltage changes can trigger action potentials in quiescent neurons [17,20]. However, even lower field magnitudes, such as those generated in the brain during transcranial electrical stimulation techniques, can still modulate spike rate and timing [21–23].

Despite knowing how electrical stimulation affects single neurons, predicting its impact on population activity is challenging due to neuronal variability in morphology, electrophysiology, and synaptic properties [24–26]. This is a crucial problem because the capacity of electrical stimulation to modulate brain function depends on its global effects on brain tissue. From a practical standpoint, this issue also presents significant challenges in designing stimulation protocols that achieve the desired outcomes. Current approaches to modulate brain activity at the population level often involve matching the stimulation frequency to the overall temporal pattern of neuronal activity (as estimated from the local field potential or EEG signals) [27,28], using predefined stimulation parameters that have been shown to effectively modulate clinical symptoms in pathological conditions [29] or, more recently adaptive/closed-loop approaches to modulate more selectively brain activity [30–32]. While valuable, these approaches must be complemented by a deeper understanding of how electricity affects brain activity.

One potential approach to achieve that is to study the effects of electrical stimulation across multiple spatial scales, ranging from single neurons to populations. Electrophysiological extracellular recordings enable recording the activity of a large number of neurons but they are susceptible to electrical artifacts during stimulation. An alternative is to use 2-photon imaging with fluorescent indicators that are sensitive to either the presence of calcium ions [33] or to changes in the membrane potential [34]. These techniques offer the potential to monitor the overall activity of entire regions using wide-field imaging, while also allowing the user to zoom in and monitor the activity of hundreds or thousands of neurons simultaneously, as well as individual neurons or specific cellular compartments [35]. Genetic tools that are readily available in mice enable these recordings to be performed in specific cell types [36], providing a valuable resource for understanding how electrical stimulation can affect different cells in the brain. Thus, this approach holds potential for the development of new stimulation paradigms that are better suited for specific applications.

A straightforward method for combining electrical stimulation with imaging techniques is to place the electrodes directly on the brain’s surface (or dura mater). However, this approach may significantly limit the field of view, relying on using multiple small electrodes and imaging in the space between them, an approach that has been taken for recording purposes [37]. In this work, we propose a different solution: the development and use of electrodes for stimulation that, by being transparent, are fully compatible with imaging techniques. The ideal electrodes for this purpose should possess high biocompatibility, transparency, and charge injection capabilities sufficient to generate electric fields in the brain that are comparable to those produced by various techniques in humans.

A material that meets these requirements is PEDOT:PSS (poly(3,4-ethylenedioxythiophene) polystyrene sulfonate). PEDOT:PSS is a conductive polymer that has been shown to be highly biocompatible and well-suited for recording electrical activity and stimulating brain tissue [38]. For instance, metal electrodes coated with PEDOT:PSS exhibit lower impedances, resulting in improved signal-to-noise ratio [39,40]. Coating metal electrodes with PEDOT:PSS also increases the amount of charge that can be stored and used for stimulation purposes, with an increasing thickness leading to higher charge injection capacity [41,42]. This feature is primarily due to the mixed ionic/electronic conductance of PEDOT:PSS material that confers volumetric capacitance to PEDOT:PSS films [43]. Recent work has also demonstrated that electrodes coated with PEDOT:PSS can induce neural responses in the mouse visual cortex for over a year, thus highlighting the high biocompatibility of this material [44].

In this study, we directly inkjet-print PEDOT:PSS to produce an electrode ideal for neurostimulation. Inkjet printing is a drop-on-demand patterning technique with micron precision [45] that allows rapid prototyping and testing while limiting wasted material. This technique is gaining popularity for producing devices that interface with biological tissue, with the goals of recording and stimulation [46]. One of the main advantages of using PEDOT:PSS inks is that their properties can be easily altered [47] to achieve films with desired electrical, mechanical properties while maintaining stable performances in water [48]. Thin films of PEDOT:PSS can be highly transparent [49,50], and PEDOT:PSS, while potentially less conductive than typical metals, does not suffer from some limitations of other commonly used transparent materials for electrodes (for a review, see [51]). For instance indium-tin-oxide (ITO), one of the most used inorganic materials for transparent electrodes, is expensive, brittle and limited in terms of flexibility [52]. While graphene is an excellent alternative for transparent electrodes [53–55], cleanroom fabrication remains less cost-effective, and research to improve inks formulations for printing is still ongoing [56,57].

Here, we demonstrate that inkjet-printed PEDOT:PSS electrodes can exhibit high charge injection capabilities while maintaining high transparency. We evaluate their transparency under 2-photon imaging conditions and perform in-vivo experiments in mice, combined with a finite-element model, to determine the maximum electric field that these electrodes can generate. Our findings indicate that the electrodes we fabricated can produce electric fields in the mouse brain up to 300 V/m, which meets the requirements of many electrical stimulation modalities. Therefore, inkjet-printed electrodes represent a promising new tool for studying the effects of electrical stimulation in combination with simultaneous imaging techniques.

## RESULTS

We fabricated a device with a transparent electrode for stimulation using a hybrid approach that combines standard microfabrication techniques in a cleanroom environment with inkjet printing (**Figure 1A**). Inkjet printing enables the design and quick patterning of potentially multi-layered materials with varying thicknesses on desired substrates. Our primary innovation in this study is to leverage this process to print conductive polymer electrodes suitable for electrical stimulation.

**Figure 1.**
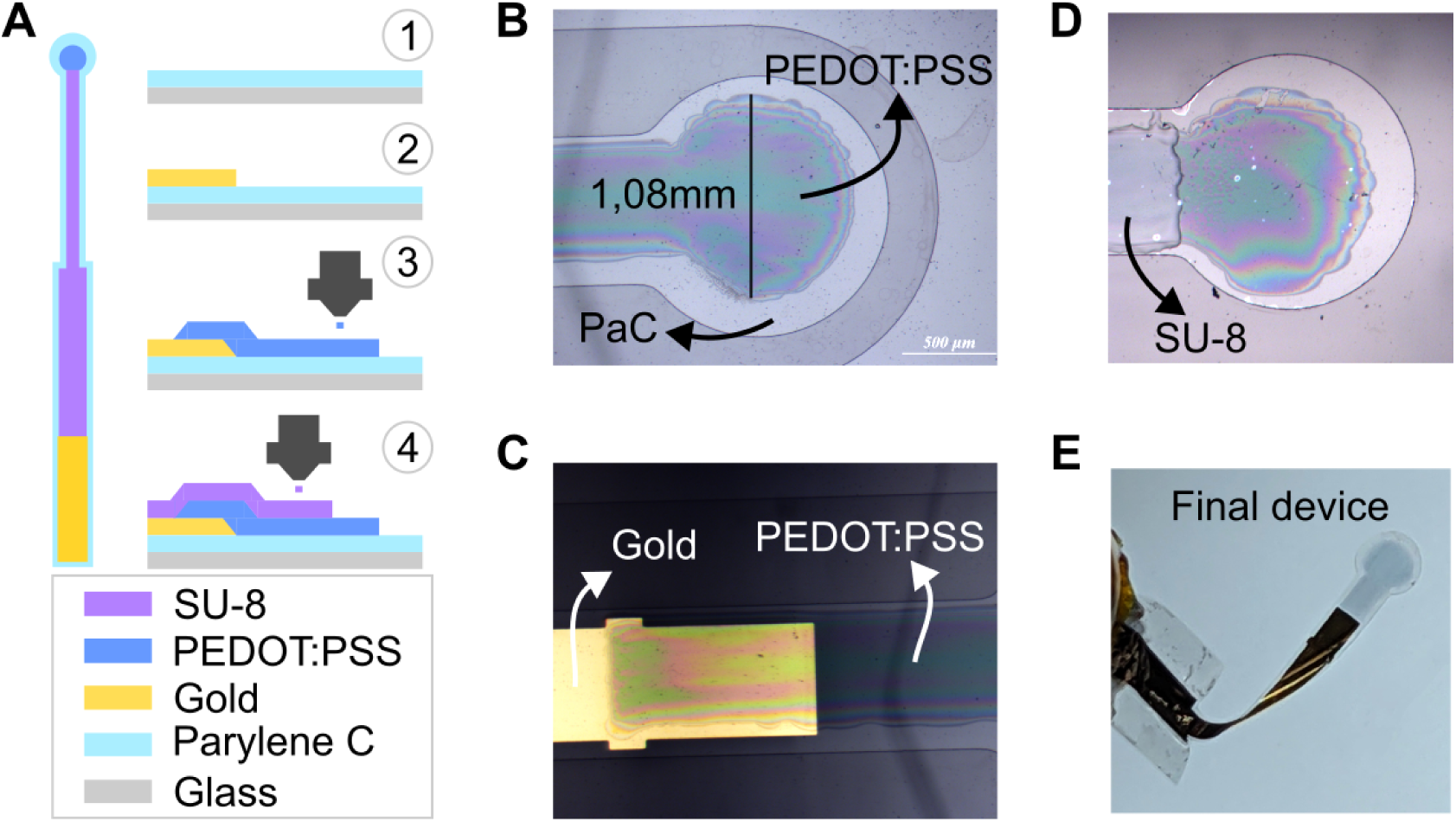
Device fabrication. **A**: Schematics of the device and the fabrication process. Vapor deposition, photolithography and gold deposition were used to deposit Parylene C and create the connection line. The electrode, a single circle with a 1 mm diameter, was inkjet printed using a PEDOT:PSS ink. Insulation was applied by printing multiple layers of the photoresist SU-8. **B**: Picture of a representative device after printing PEDOT:PSS. **C**: Interconnection between gold and the printed PEDOT:PSS layer. **D**: Picture of a representative device after printing 3 layers of SU-8. **E**: Example of released device ready for testing.

Briefly, the fabrication process began with the chemical vapor deposition of Parylene C (PaC) on a glass substrate, to provide a layer for encapsulation and electrical isolation. Next, photolithography was used to pattern the device-to-generator connection, followed by the deposition of 120 nm of gold using thermal evaporation. After defining the device’s outline via photolithography and etching (to facilitate the release from the substrate), we then inkjet printed PEDOT:PSS, using a custom ink formulation [48] to define the electrode. The electrode consists of a single circle with 1 mm diameter (**Figure 1B**, see *Discussion* for an explanation on the chosen size) and the ink was overlapped with the gold lines to create an electrical connection between the two (**Figure 1C**). A SU-8 ink was then printed to isolate the device leaving only the exposed electrode (**Figure 1D**). The device was subsequently removed from the substrate, resulting in a highly transparent electrode within a highly flexible device (**Figure 1E**). A more detailed explanation of the fabrication process is reported in the *Materials and Methods* as well as a full schematics of the process in **Supplementary Figure 1**. To explore the importance of thickness, we fabricated devices with varying numbers of printed PEDOT:PSS layers, thus controlling this parameter. This allowed us to estimate the electrical and optical properties across different devices.

### Electrical characterization

The overall thickness of the device was determined by the number of printed PEDOT:PSS layers (**Figure 2A**). Electrodes made from a single layer had a thickness of 350 ± 63 nm (mean ± SD, n = 5 electrodes) with each additional layer further increasing the total thickness. For example, electrodes with four layers of PEDOT:PSS exceed 1 μm in thickness. While the volumetric capacitance of PEDOT:PSS suggests that adding more layers should enhance the maximum current injection capacity of electrodes, this increased thickness may compromise the device’s transparency, potentially limiting its use in imaging applications. Here we will first characterize the electrical properties of the electrodes, followed by an analysis of their transparency in the subsequent section.

**Figure 2.**
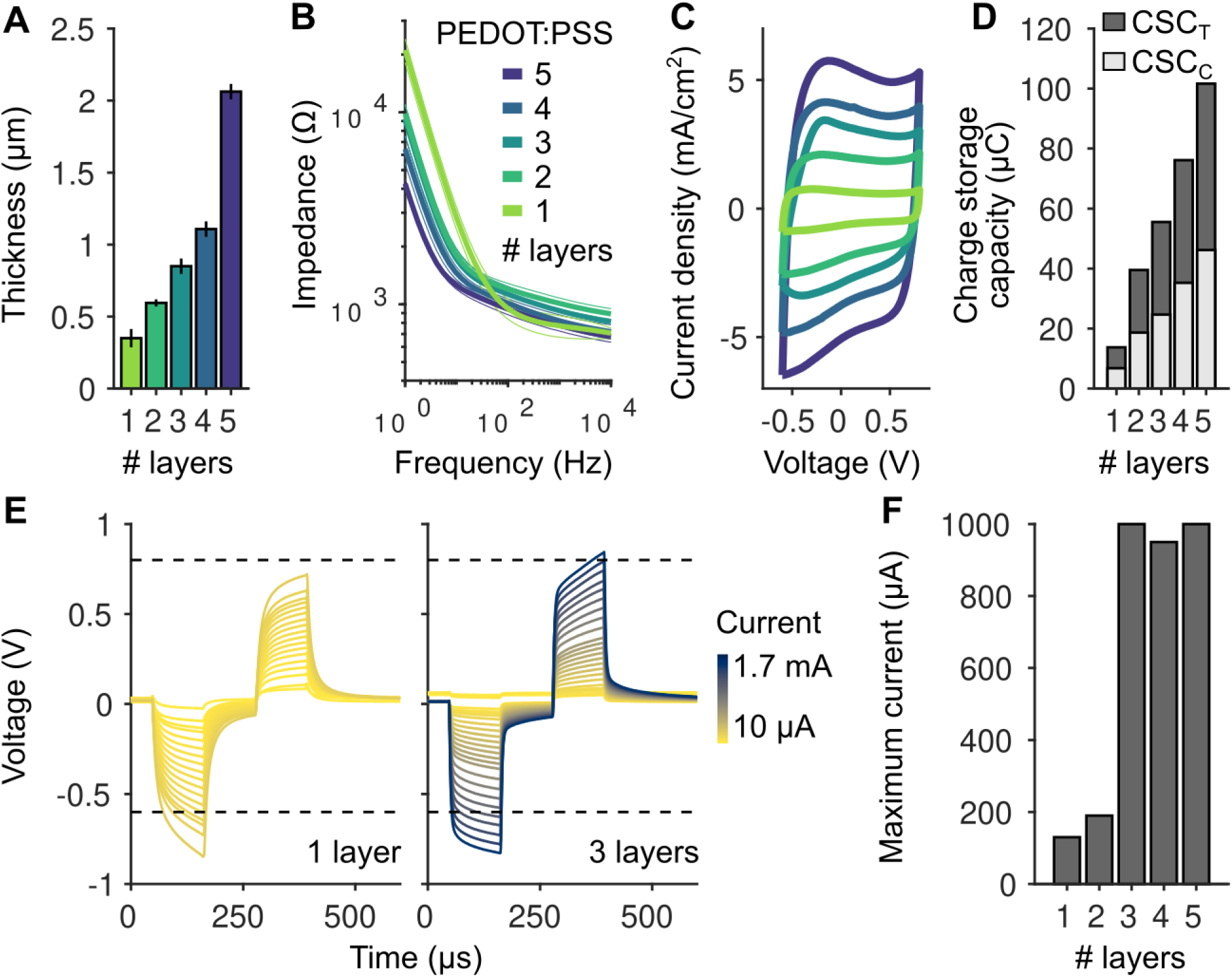
Electrical properties of inkjet printed electrodes. **A**: Thickness of electrodes made of a different number of printed PEDOT:PSS layers (n = 5, 3, 5, 2, 2 electrodes for 1-5 layers respectively). **B**: Electrical impedance spectroscopy measurements for electrodes made of different numbers of inkjet printed PEDOT:PSS layers. Thick lines represent mean values across electrodes and thin lines the error bars estimated as standard deviation (n = 9 electrodes for 1 printed layer and and n = 3 for the others). **C**: Cyclic voltammetry measurements for electrodes made of different numbers of inkjet printed PEDOT:PSS layers. **D**: Charge storage capacity (cathodic: light gray, total: dark gray) estimated from the cyclic voltammetry measurements for different numbers of layers. **E**: Example traces (for one and three layers PEDOT:PSS) from pulse tests to estimate the maximum current that can be applied using the electrodes. **F**: Maximum current that can be injected using the electrodes before reaching the water window ([−0.6 0.8] V).

We characterized the electrical properties of the devices in phosphate-buffered saline (PBS). We first estimated the sheet resistance of printed PEDOT:PSS using four-point probe measurements (see *Materials and Methods*). We performed the measurements using both one layer and two layers. We estimated values of 346 ± 3 Ω/◻ and 174 ± 6 Ω/◻ (“ohms per square”, mean ± SD, n = 7 measurements for one printed layer and n = 6 for two layers), corresponding to conductivity values of 131 ± 1 S/cm and 133 ± 5 S/cm, consistent with values obtained with the same ink formulation [48]. Note that the similar conductivity (their values are not significantly different, p = 0.43, t-test) is expected considering that the prints have different thickness (see next section) but are made of the same material. Nevertheless, similar values confirm that there is no electrical barrier between two layers.

We next performed Electrochemical Impedance Spectroscopy (EIS) to determine the overall impedance of the electrodes as a function of the number of printed layers (**Figure 2B**). We found that the low-frequency impedance decreased with increasing number of printed layers while staying lower than 1 kΩ for frequencies higher than 1 kHz. At the lowest frequency tested (1 Hz), the impedance for one printed layer was just 21 ± 4 kΩ (mean ± SD, n = 9 electrodes), a low value that allows to use this electrode with a large set of commercially available stimulators.

We then estimated the amount of charge that the electrodes could store as a function of the number of layers using Cyclic Voltammetry (CV, **Figure 2C**). The CV, representing the current density over the potential, has approximately a rectangular shape. The absence of noticeable redox peaks implies no significant electrochemical reactions and instead indicates a double-layer charging. Charge storage capacity (CSC), which is the total charge transferred during the cyclic sweep, can be determined directly from the CV by integrating the current over the potential range (−0.6 to 0.8 V). Increasing the number of printed layers, and the resulting increased thickness of the electrode, leads to higher current storage capacity (**Figure 2D**), as expected considering the increased capacitance of the device.

Finally, we applied test pulses (**Figure 2E**) to identify current values that lead to voltages that cross the water window for PEDOT:PSS vs Ag/AgCl ([−0.6 0.8] V) either in the anodic of cathodic phases. We found a value of max 130 μA for one layer electrodes before crossing the water window. This value increases to 190 μA for two layers and jumps abruptly to about 1 mA at higher numbers of layers (**Figure 2F**).

In summary, the results of our electrical characterization suggest that the electrical properties of the circle electrode can be tuned by altering the number of printed layers. In general, the impedance and maximum current that can be safely injected suggest that already one single layer of PEDOT:PSS can be used to inject a significant amount of current and the low impedance allows its use with regular commercial stimulators. Increasing the number or layers leads to higher maximum currents but the increased thickness of the device can become problematic because it leads to lower transparencies, a topic that we address in the following section.

### Transmittance measurements

We aimed to estimate the transparency of our devices using two distinct methods (**Figure 3A**). First, we employed a 1-photon spectrophotometer to measure the transmittance of multiple PEDOT:PSS printed layers. This method quantifies the proportion of light passing through the material relative to the initial incident light intensity (**Figure 3A**, left). Second, we sought to estimate transmittance under 2-photon excitation conditions (**Figure 3A**, right). In this setup, a pulsed near-infrared laser was used to excite molecules in the sample by simultaneous absorption of two photons, and the resulting fluorescent emission light was collected. The overall transmittance can then be estimated by comparing the measured fluorescence signal in presence of the electrode to that obtained without the electrode. As both the excitation and emission light are attenuated by the electrode in the 2-photon setup, we anticipated lower transmittance values compared to the spectrophotometer measurements. Importantly, the spectrophotometer-derived transmittance values can be directly used to predict the transmittance under 2-photon conditions (see *Materials and Methods*). This allows for the generation of theoretical predictions, which we could test with a 2-photon microscope.

**Figure 3.**
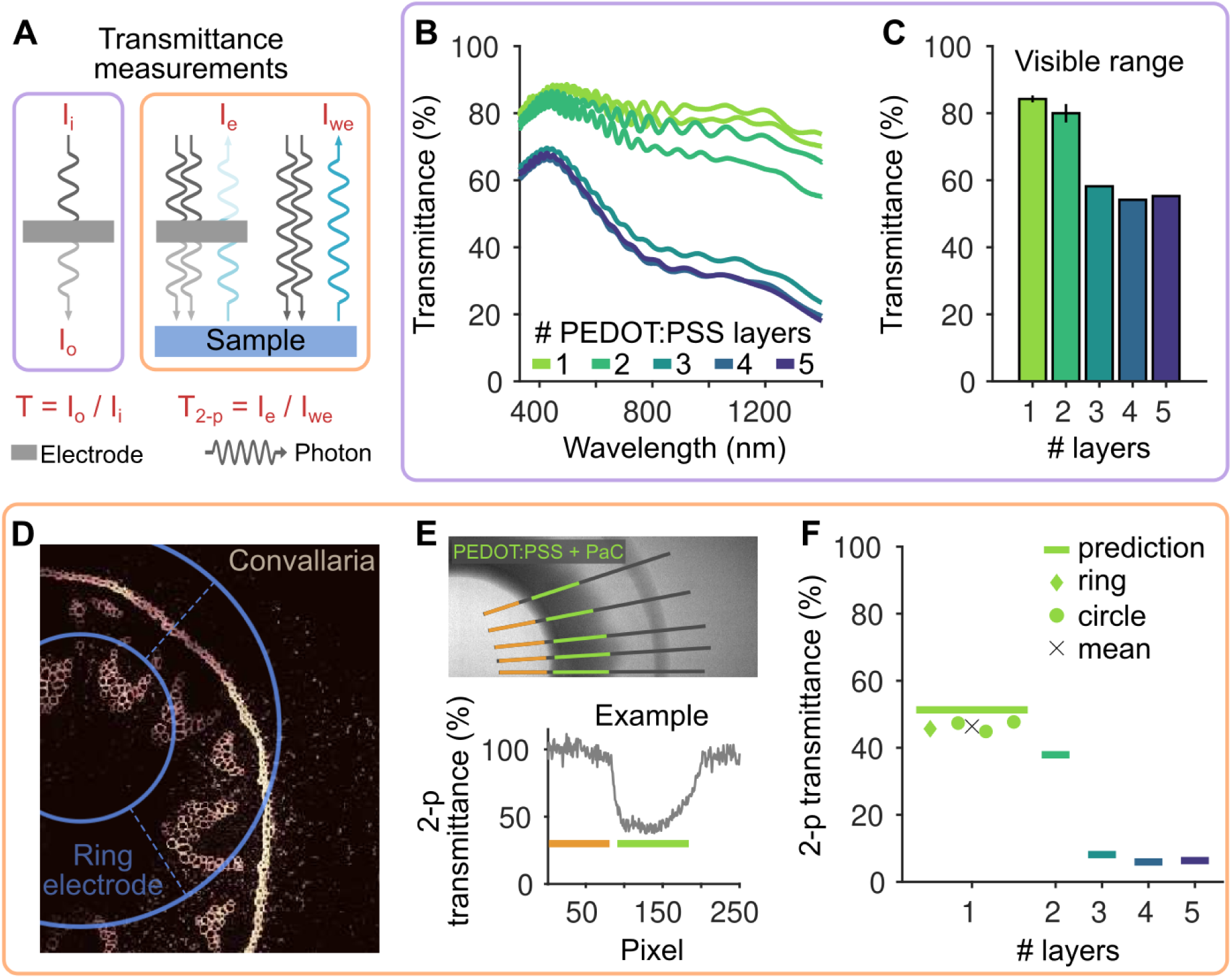
Optical properties of inkjet printed electrodes. **A**: Schematics of the tests performed to estimate the transmittance of the electrodes. One set of tests relied on measurements with a spectrophotometer (violet color, relative to panels B and C), which estimates how much light passes through the sample compared to the initial one (*I_i_*: input light, *I_o_*: output light). The other set of tests are in 2-photon conditions (orange color, relative to panels D-F), where two photons stimulate a sample and the following emission light is collected by the sensor. Transmittance can be estimated by comparing the emission light collected through the electrode (*I_e_*) and without the electrode in the light path (*I_we_*). **B**: Light transmittance as a function of light wavelength for an increasing number of printed PEDOT:PSS layers. **C**: Average transmittance in the visible range (380-750 nm) estimated from the data in B. Errorbar indicates mean and STD (n = 2, 2, 1, 1, 1 electrodes for 1-5 printed layers respectively). **D**: 2-photon imaging of a glass slide containing the fluorescent plant Convallaria majalis. A ring-like electrode made of a single layer of PEDOT:PSS is placed on the top of the slide (the position is drawn in blue). **E**: 2-photon experiment to estimate the transparency of electrodes made of a single printed layer of PEDOT:PSS. (Top) A ring electrode is placed on the top of a fluorescent glass slide. Manually selected lines for the analysis are indicated in gray while green indicate the location of the PEDOT:PSS + PaC electrode. Orange lines represent the baseline considered, where no electrode is present. (Bottom) Example of the estimated transmittance across one selected line (after correcting for the background). **F**: Estimated transparency in 2-photon imaging conditions of electrodes made of a single printed layer of PEDOT:PSS. Symbols represent the experimental measurements and black cross the mean values across electrodes. The solid lines represent the theoretically predicted transmittances derived from the data in panel B (see *Materials and Methods*).

We first estimated the transmittance by printing 2×2 cm squares with varying numbers of PEDOT:PSS layers (1 to 5) and analyzing the amount of light passing through the material. These measurements were performed at distinct wavelengths using a one-photon stimulation spectrophotometer (**Figure 3B**; see *Materials and Methods*). Approximately 84% of light in the visible range (380–750 nm) passes through a single layer of PEDOT:PSS (**Figure 3C**). Two printed layers of PEDOT:PSS exhibit an 80% transmittance, while further increasing the number of layers results in a rapid decrease in transmittance.

We then conducted tests under 2-photon imaging conditions. Initially, we placed a single-layer PEDOT:PSS electrode (ring-shaped to fit within a single field of view) on top of a glass slide containing a fluorescent plant (Convallaria majalis) to determine if fluorescent structures could be visualized beneath the electrode. Our results confirmed this visualization was possible (**Figure 3D**). To quantitatively estimate the transmittance of the electrodes, we imaged uniform fluorescent slides, which allowed us to directly quantify the emission light collected beneath the electrode compared to a region without the electrode in the light path. We extracted the light intensity over multiple manually drawn lines, identified the electrode location, and estimated the light loss under the electrode compared to a location without the electrode, after correcting for the background (**Figure 3E**; see also *Materials and Methods*). In the represented example, the transmittance of a single layer of PEDOT:PSS was just below 50%. This estimate was confirmed using multiple full-circle electrodes (47 ± 2 %, mean ± SD across 3 electrodes, **Figure 3F**).

Based on the transmittance values obtained using the spectrophotometer, we computed the theoretical transmittance expected under 2-photon conditions (lines in **Figure 3F**), accounting for the multiple obstructions of the light path by the electrode (see *Materials and Methods*). The experimental values were slightly below the theoretical predictions, suggesting that spectrophotometer measurements alone can be used to derive accurate estimates of transmittance in 2-photon experiments. Notably, a measured transmittance of about 50% using a spectrophotometer, as measured for 3 to 5 layers, corresponds to only 5-10% transparency in 2-photon conditions, making it less suitable for imaging experiments. However, a two-layer electrode may only lose approximately 10% transmittance compared to a single-layer one.

### In-vivo experiments and computational modeling

Considering that a single-layer PEDOT:PSS electrode can deliver currents exceeding 100 μA (**Figure 2F**) while preserving approximately 50% transmittance for 2-photon imaging (**Figure 3F**), we opted to use a single PEDOT:PSS printed layer for subsequent in vivo experiments, where we aimed to directly estimate the electric field generated in the brain by the electrodes.

The effects of electricity on brain activity depend on the electric field generated within the tissue [12]. Thus, we aimed to estimate the magnitude of the electric field produced by a single-layer PEDOT:PSS electrode placed on the dura mater of an anesthetized mouse. Estimating the electric field can be achieved by measuring voltage fluctuations induced by the stimulation at various locations within the brain. Thus, we recorded voltages in the mouse brain using rigid electrodes referenced to a screw placed in the back of the animal skull. For stimulation, we placed the transparent PEDOT:PSS electrode on the mouse dura and a metal screw counter electrode in the front part of the brain, to approximate a monopolar stimulation configuration (**Figure 4A**). We applied sinusoidal 10 Hz current–controlled stimulation at multiple amplitudes across multiple repetitions. We limited the maximum current to 100 μA, just below the maximum value that was identified during the device characterization, as higher currents resulted in the electrode losing functionality (see **Supplementary Figure 2**). The electrode was lowered perpendicularly to the brain surface, and voltage fluctuations induced by the stimulation were recorded at multiple depths (**Figure 4B**). We estimated the electric field component perpendicular to the brain surface by calculating the difference between the amplitudes of the measured sinusoidal voltages at consecutive depths and dividing by their distance. We found that the electric field magnitude scaled linearly with the applied current (slope of the fit, β = 0.999, R² = 0.9998; top plot in **Figure 4C**). At an applied current of 100 μA, the peak electric field was approximately 30 V/m (**Figure 4C**). This field strength could be sufficient to elicit action potentials in some neurons [17], as the minimum activation threshold is estimated to be around this value (see *Discussion*). Additionally, a measurement of 30 V/m approximately 1 mm from the center of the stimulating electrode suggests that the electric field directly beneath the electrode may be an order of magnitude larger.

**Figure 4.**
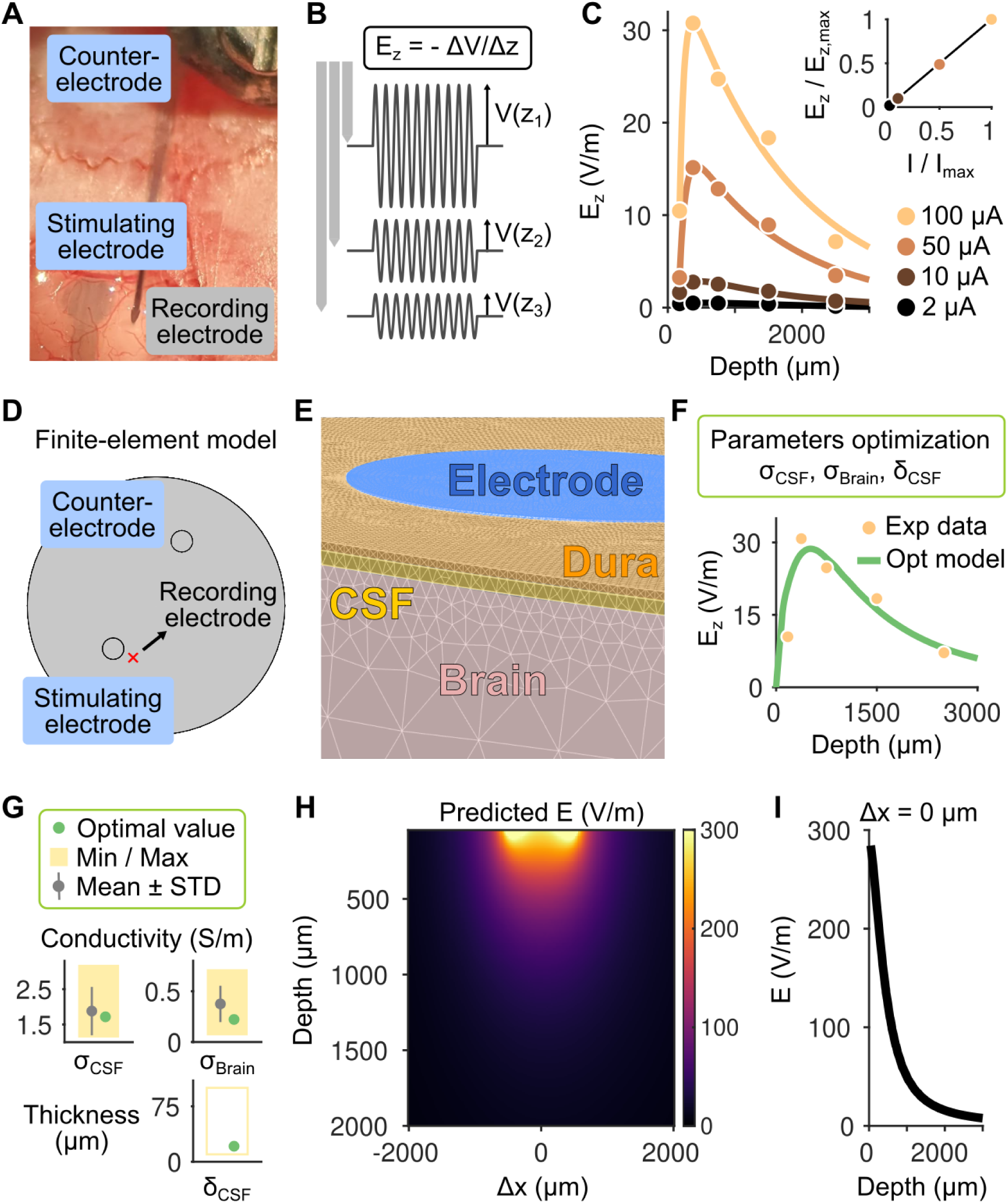
Combined in-vivo experiment and computational modeling to estimate the electric field generated by the inkjet-printed electrode. **A**: The flexible printed PEDOT:PSS electrode is placed on the mouse brain for stimulation, together with a screw counter electrode in the front. A rigid electrode is used to record voltage fluctuations due to a sinusoidal current stimulation (10 Hz). **B**: The component of the electric field perpendicular to the brain surface (E_z_) is estimated by taking the difference of the amplitude of the recorded sinusoidal voltages at multiple depths (schematic representation). **C**: Estimated electric field (E_z_) for four different currents applied, indicated by different colors. Lines indicate a double exponential fit for each stimulation amplitude. The top plot shows the linearity between current applied and the electric field (both of which are normalized by the values corresponding to the maximum current applied). The black line is the identity line. **D**: Geometry used for the FEM to simulate the experimental conditions. The stimulating and counter electrodes match the geometry and location of the experiment. **E**: Representation of the FEM meshing with different colors representing different tissue types (electrode, dura, CSF, brain). **F**: Experimental data and simulation results for a model in which three parameters (the conductivities of the brain and CSF and the thickness of the CSF layer) were optimized. **G**: Comparison of the optimal parameters identified with the means and standard deviations derived from multiple studies and reported by Sim4Life (see *Materials and Methods*). Yellow rectangles represent the minimum and maximum values reported in the literature, which were used as the bounds for the optimization procedure. The CSF thickness varies depending on the location in the brain and therefore we left quite large boundaries for the optimization. **H**: Predicted electric field beneath and close by the stimulating electrode. The 2D distribution represents the electric field magnitude derived from the model that best matches the experimental data. **I**: Dependence of the electric field magnitude on the depth directly below the center of the stimulation electrode.

To test this hypothesis, given the practical challenges of measuring voltage fluctuations directly beneath the electrode, we developed a computational approach where we numerically simulated our experimental conditions while optimizing the model parameters based on the experimental data. By doing so, we aimed to estimate the electric field distribution in regions where direct measurements were unfeasible. Specifically, we implemented a finite-element model (FEM) where we positioned the active electrode and counter-electrode as in the experimental setting and assigned them the same geometrical properties as in the experiments (**Figure 4D**, see *Materials and Methods*), an approach that is standard to simulate electrical brain stimulation [58–60]. We modelled the electric currents reaching the brain, passing through the dura mater and the cerebrospinal fluid (CSF) underneath, three tissue types that we defined in our simulation (**Figure 4E**). We applied 100 μA current to the simulated PEDOT:PSS electrode (as in the experiments) and set the counter electrode to ground to close the circuit. Rather than manually assigning electrical conductivities to the tissues, an optimization algorithm was used to determine the optimal set of parameters that best matched the experimental data (**Figure 4F**). This optimization process focused on the conductivities of the CSF and brain tissue, as well as the thickness of the CSF layer (the subarachnoid space), aiming to minimize the difference between the simulated and experimentally measured perpendicular component of the electric field at the corresponding location (see *Materials and Methods*). We found a set of parameters that led to an electric field that closely matched the data in the corresponding position (**Figure 4F**). We further checked whether the optimal parameters were consistent with the ones commonly used in the literature (see *Materials and Methods*) and found them to be within one standard deviation from the mean (**Figure 4G**), suggesting that our experimental measurements aligned with predictions from basic current flow across the tissues. The optimal value for the thickness of the CSF layer was also compatible with experimental data from previous reports [61]. We then analyzed the distribution of the electric field magnitude generated by the model in a full 2D space that included areas directly beneath the stimulating electrode (**Figure 4H**). We found that the electric field directly below the center point of the electrode reached approximately 300 V/m (**Figure 4I**), a value about 10 times the minimum threshold for direct neuronal activation [62].

In summary, by combining an in-vivo experiment with a parameters-optimized FEM we were able to estimate the full distribution of the electric field inside the brain, including directly beneath the stimulating PEDOT:PSS electrode. This electric field distribution and magnitude could modulate large neuronal populations while minimizing variability in stimulation effects often associated with non-uniform field distributions. This advantage is particularly significant compared to multi-electrode stimulation approaches, which often generate highly non-uniform electric fields.

## DISCUSSION

In this work, we showed that a ∼1 mm diameter electrode made of a single PEDOT:PSS printed layer (thickness ∼350 nm) allows a transmittance of approximately 50% in 2-photon imaging settings and can inject currents of up to ∼130 μA, which correspond to an electric field magnitude of maximum 300 V/m in the mouse brain when delivered on the dura mater. These results were derived combining microfabrication, inkjet printing, electrical and optical device characterization, in-vivo experiments and computational modeling with parameters optimization procedures.

Our strategy for the fabrication of the device was hybrid combining standard cleanroom techniques with inkjet printing. This approach was chosen to facilitate the analysis of the printed part of the device without adding the complexity of potential issues that may arise from the connector part, as our cleanroom fabrication procedure is already well established. Future work will focus on developing fully printable devices [46], for instance by directly patterning the connectors with gold ink. This approach holds significant promise for advancing the development of flexible devices [63], by leveraging the cost-effective fabrication provided by printing techniques, an additive manufacturing method that minimizes waste and can be performed without a cleanroom environment.

A single, relatively large electrode design was chosen for two primary reasons. First, this configuration served as a proof-of-principle demonstration that inkjet-printed electrodes of this type are viable for neural stimulation. Second, minimizing electric field variations beneath the electrode is challenging with multi-electrode arrays. The large, single electrode design mitigates this challenge, enabling more uniform and controlled neuronal stimulation, a feature that is further enhanced by our direct estimation of the electric field generated by the electrode. Nevertheless, the fabrication method, using inkjet printing with a previously reported resolution of approximately 30 μm [48], allows for future development of multi-channel devices, if necessary, using the same approach.

Our tests in PBS showed that adding printed layers of PEDOT:PSS increases the maximum charge that can be injected. However, this increase in charge injection capacity correlated with a decrease in electrode transparency. To achieve higher current densities in applications where transparency is critical, several strategies can be considered. Modifications to the ink formulation [64], including alternative dopants or functionalization strategies [65], may improve charge injection without compromising transparency. Multi-layer coatings, for instance using silver nanowires/PEDOT:PSS films, represent another promising approach to enhance charge injection while maintaining or even improving transparency [66]. Future studies should also address the challenge of reliable current delivery at elevated levels to mitigate delamination at the gold/PEDOT:PSS interface. Strategies to achieve effective current delivery at elevated levels may include surface treatments to enhance adhesion [67,68] or the incorporation of a buffer layer to reduce stress.

By combining in-vivo experiments with a parameter-optimized FEM approach, we were able to estimate the electric field directly below the electrode. Finding that the optimal parameters for the fit are consistent with values reported in the literature validated our modeling approach and the subsequent use of the model to estimate the electric field directly below the electrode. While direct measurements could be performed to test the predictions, they would require insertion of electrodes at extreme angles to measure the most superficial layers of the cortex. Etching a small hole to allow for the insertion of a recording electrode directly below the stimulating PEDOT:PSS electrode would also greatly alter current flow and thus yield an altered estimate of the electric field. Nevertheless, the close agreement between the experimental data and the simulation results, obtained using physiologically plausible parameter values compatible with the literature, indicates that the fundamental principles of current propagation adequately describe the observed experimental outcomes. This also confirms that this type of modeling predicts the actual measured electric fields in the brain with great accuracy [10].

Our results suggest that the electric field achieved under the electrode may be as high as 300 V/m. This raises the question of how this number compares to common applications of electrical brain stimulation. Different electrical brain stimulation modalities generate electric fields that span different orders of magnitudes. Invasive techniques, such as deep brain stimulation (DBS) or cortical microstimulation generate electric fields from hundreds to thousands of V/m depending on electrode size and distance from the electrode [58,69]. Electroporation techniques rely on electric fields in the range of 10^4^ to 10^7^ V/m, depending on the desired effects [70]. On the other hand, transcranial techniques, such as transcranial direct and alternating current stimulation (tDCS, tACS), generate electric fields at the cortical level of max about 1 V/m [10,59], although electroconvulsive therapy can generate fields (of short duration) of up to hundreds of V/m [71]. Temporal interference stimulation, another form of transcranial electrical stimulation, can generate electric fields of maximum 0.4 V/m in deeper areas in the brain [72,73]. Electric fields in the order of 100 V/m in the human cortex, generated using transcranial magnetic stimulation, can also induce motor responses [74,75]. In general, a threshold of 15-20 V/m is considered as the minimum to induce action potentials and 5 times that value is considered enough for robust activation [17,62].

Considering this evidence from the literature, our electrodes are well suited to study the effects of different electrical stimulation techniques on brain activity, to either induce spiking activity or modulate neuronal activity with lower amplitude electric fields, which are nevertheless effective in modulating neuronal activity [13,76]. For instance, while the effects of low-amplitude electrical stimulation on firing activity have been characterized for decades in-vitro and in-vivo using extracellular recordings [14,21–23,77–83] or calcium imaging [84–88], a direct estimation of the membrane voltage fluctuations induced by electrical stimulation have been mainly limited to somatic compartments [15,17,77,89,90] or estimated only in in-vitro preparations [91]. With the rapid development of new indicators to directly monitor changes in membrane voltage in mice [34,92,93], our transparent electrode represents the ideal tool to test predictions on the direct effects of electrical stimulation on the neuronal membrane potential from decades of computational modeling work, including those from more recent years [18,94–96].

While rigorous testing is crucial to ensure long-term biocompatibility and functionality [44], our device demonstrates a significant breakthrough by achieving both cost-effectiveness in development and rapid testability. This achievement holds far-reaching implications, extending beyond research applications to clinical settings that demand both adequate stimulation performance and optical access.

## MATERIALS AND METHODS

### Device fabrication

The device was made using a combination of standard cleanroom techniques and inkjet printing and the detailed process is represented in **Supplementary Figure 1**.

Glass slides, which were used as the substrate, were sonicated in a 2% soap solution, then rinsed, and sonicated in acetone and isopropanol. A 3 µm layer of Parylene C (PaC) was deposited via chemical vapor deposition *(Specialty Coating Systems, PDS 2010, USA)*. Photolithography with AZ nLOF 2070 *(Microchemicals, Germany)* negative photoresist was employed to pattern the gold structures. This involved spin-coating (500 rpm for 10 seconds, and 3000 rpm for 40 seconds), soft baking (2 minutes at 110 °C), exposure (186 mJ/cm^2^ at the i-line wavelength), post-exposure baking (2 minutes at 110 °C), and development in AZ 826 MIF developer. Titanium (10 nm) and gold (120 nm) layers were deposited via electron beam and thermal evaporation respectively, followed by a lift-off process involving acetone and isopropanol. A second photolithography step with AZ 10XT *(Microchemicals, Germany)* positive photoresist defined the device’s outline. After spin-coating (3500 rpm for 35 seconds), soft baking (2 minutes at 110 °C), exposure (480 mJ/cm^2^ at the h-line wavelength), and development in AZ developer, reactive ion etching (RIE) *(Oxford instruments, Plasmalab 80+ RIE, UK)* with O_2_ and CF_4_ gasses was employed to etch the PaC to define the outline.

A piezoelectric inkjet printer *(DMP 2800, Fujifilm Dimatix, Santa oven Clara, CA, USA)* patterned the PEDOT:PSS electrodes and the insulating SU8 layer. Custom-formulated PEDOT:PSS ink was used based on previous work [48]. It includes commercially available PEDOT:PSS water dispersion *(Clevios PH1000 by Heraeus)*, DMSO *(Thermofisher, USA)* (10% w/w), IPA (5% w/w), TWEEN 20 (0.5% w/w), and GOPS (1% w/w). The ink was mixed, sonicated, filtered, and printed with precise alignment. Printed layers were cured, and excess material was washed off. Samba cartridges with a 2.4 pL drop volume and 15 µm spacing patterned the PEDOT:PSS electrodes on the gold lines, then the prints were cured at 130 °C for 10 minutes. SU-8 2002 *(SU-8 2002; MicroChem Corp.)* was used for insulation due to its excellent chemical and mechanical properties. It was diluted with cyclopentanone for better printability and applied in three layers, followed by drying (95°C for 60 seconds) and UV cross-linking. The devices were finally released from the glass substrate by immersion in deionized water.

### Device electrical characterization

Sheet resistance was estimated using four-point probe measurements with a Jandel RM3000 unit (*Jandel, UK*). Electrochemical impedance spectroscopy (EIS) was performed using a PalmSens4 potentiostat (*PalmSense BV, The Netherlands*) to analyze the electrical properties of PEDOT:PSS. Measurements were taken across frequencies from 1 to 10^6^ Hz in a two electrode setup with phosphate buffered saline (PBS). PEDOT:PSS served as the working electrode, and silver-silver chloride (Ag/AgCl) was used as the counter and reference electrodes. A sinusoidal potential with a 10 mV amplitude was applied to the WE. The impedance spectrum exhibited capacitive behavior at lower frequencies and increasingly resistive behavior at higher frequencies. Increasing the number of PEDOT:PSS layers resulted in a noticeable decrease in impedance magnitude as seen in **Figure 2B**. Cyclic Voltammetry (CV) was used to study the redox behavior of PEDOT:PSS in PBS by linearly sweeping the potential of the working electrode between −0.6 to 0.8 V at a scan rate of 1 V/s with steps of 1 mV. The cyclic voltammograms indicate no significant electrochemical reactions and thicker layers show larger currents, indicating enhanced capacitance (**Figure 2C**). The charge storage capacity (CSC), representing the total charge transferred during cyclic sweeps, was determined by integrating the current over the potential range (−0.6 to 0.8 V) from the cyclic voltammetry (CV) results. Increasing the thickness of PEDOT:PSS layers enhance the surface area available for charge storage, thereby increasing CSC as seen in **Figure 2D**. Charge Injection Capacity (CIC) was determined by applying biphasic current pulses (100 µs duration) *(Metrohm Autolab, Nova 2.1)* to electrodes made of different PEDOT:PSS layers and identifying the maximum current that could be used before crossing the water window (**Figures 2E,F**).

### Device transparency characterization

The transparency of the PEDOT:PSS electrodes was evaluated by assessing the transmittance of light through a printed square (2×2 cm) with various numbers of printed layers (1 to 5), using a spectrophotometer (*UV-2600, Shimadzu, Japan*). The instrument is equipped with a light source covering the specified wavelength *w*, ranging from 300 to 1400 nm. A monochromatic light passes through the sample (the printed squares), and the spectrophotometer measures the transmittance, which is the ratio of the intensity of the transmitted light to the intensity of the incident light. This measured transmittance T(*w*) was used to predict the effective transparency for 2-photon experiments. By knowing the transmittance for the excitation laser (T_exc_) and sample emission (T_e_), we can provide an upper bound to the effective transmittance, T_2p_ = T_exc_^2^ · T_e_. The presence of the square factor is attributed to the two-photon excitation process. As the sample is excited by two photons, the excitation efficiency when crossing the electrode is reduced by a factor of T_exc_ for each photon, resulting in an overall damping factor of T_exc_^2^. The wavelength of the stimulation light of the 2-photon microscope we used is 920 nm wavelength while we collect light at 520 nm. Thus, the transmittance is expected to be T_2-p_ = T(920 nm)^2^ · T(520 nm). Considering that electrodes also contains a layer of PaC, whose transmittance we can assume to be uniform across visible light wavelengths and to be T_Pac_ = 0.97, the final transparency of the device should be no larger than T_2-p_ = T(920 nm)^2^ · T(520 nm) · T_PaC_ ^3^. In **Figure 3F**, we show the expected transparency for different layers of PEDOT:PSS deposited on PaC.

Tests in 2-photon imaging conditions were performed using a 2-photon microscope (*Ultima 2P Plus, Bruker Corporation, MA, USA*). We acquired two images: one with the electrode in the field of view and one without. Using the image with the electrode, we extracted the intensity profile of five manually defined lines. These lines crossed the electrode and included a baseline region where there was no electrode (see an example in **Figure 3E**, top). After matching the baseline intensities of the signals extracted from the two images, we divided the signals from the image with the electrode by the signals from the image without. This normalized for the non-uniform background. As a result, we obtained a baseline defined at 100%. By measuring the average signal in the area where the electrode was present, we could directly estimate the transmittance in 2-photon conditions (see an example in **Figure 3E**, bottom). Overall transmittance for each tested electrode was defined as the average across the multiple lines we defined. The same analysis was performed for either ring-like or full circle electrodes. For one electrode, we could not image the full electrode within the microscope field of view and therefore we first compared the light loss from the PEDOT:PSS electrode to the PaC and then, using a separate image, from the PaC to an area with no device. We performed this two-step procedure using exactly the same approach described above.

### In-vivo electrophysiology

All animal experimental procedures were approved and performed in accordance with the French Ministry of Higher Education, Research and Innovation (approval reference number APAFIS#22182–2019091818381938). Analgesic was administered intraperitoneal before starting the procedure (buprenorphine 0.3 mg/ml, diluted 1:10). Mice were then anesthetized with isoflurane in an induction box (4% volume in O_2_) and subsequently placed in a stereotaxic frame (*Kopf Instruments*) where the anesthesia was maintained (1–1.5%). Eyes were covered with ointment and body temperature was controlled using a heating pad. After injecting lidocaine (20 mg/ml) subcutaneously and cleaning the skin with betadine, a midline incision was performed to expose and clean the skull. Two craniotomies were performed to insert two screws to use as a counter electrode for stimulation and a reference electrode for the recording system. A large craniotomy (3 mm in diameter) was performed between visual and somatosensory cortex and the flexible electrode was placed on the top of the dura mater. A small incision of the dura was performed to insert a rigid tungsten electrode insulated with PaC (#573220, *A-M Systems, WA, USA*) approximately 1 mm from the center of the stimulating electrode. Voltage signals were filtered and amplified (model 3000, *A-M Systems, WA, USA*) before being digitized at 1 kHz (NI DAQ USB-6001, *National Instruments, TX, USA*). The same DAQ was also used to output a 1 s, 10 Hz sinusoidal voltage that was converted to current by a linear stimulus isolator (*Soterix Medical Inc., NJ, USA*). The software to control the stimulus waveform and record voltage traces was written in MATLAB (*MathWorks Inc., MA, USA*). Recordings were performed at multiple depths and for multiple trials to improve the estimation of the voltages inside the brain. After the procedure, deeply anesthetized animals were sacrificed using cervical dislocation.

### Electric field estimation

To estimate the electric field the voltage signals were filtered (low-pass Butterworth filter, 20 Hz passband and 40 Hz stopband frequencies) and the peaks and troughs of the signal were identified and used to estimate the amplitude of the recorded sinusoids. By taking the median over the multiple stimulation cycles and over the multiple trials (20 repetitions) we were able to obtain a robust estimate of the amplitude of the recorded sinusoid. The normal component (perpendicular to the brain surface) of the electric field was computed by taking the opposite of the voltage differences across consecutive recording depths and dividing by their distance. The electric field values were spatially assigned to the midpoint between consecutive recording positions. To fit the electric field profile, we minimize the sum of the residuals between a double exponential function and the experimental points.

### Finite element modeling and model fit to the data

We simulated our experimental configuration by setting up a FEM using COMSOL (*COMSOL Inc., MA, USA*). Active and counter electrodes were placed at distances approximately as in the experiments. We simulated the brain as a cylinder with a layer of cerebrospinal fluid (CSF, the subarachnoid space) and dura mater above it. We simulated the PEDOT:PSS as a cylinder of 1 mm diameter and a thickness of 500 nm, while the counter as a cylinder with the same diameter but 200 μm thick. We set the conductance of the PEDOT:PSS to be 10^4^ S/m (consistent to our estimation during the device characterization) while the counter electrode, made of stainless steel to be 1.5·10^6^ S/m. As in the experiments we applied a 100 μA inward current to the PEDOT:PSS electrode and set the counter electrode as ground. The electrodes were placed on the top of the dura mater to which we assigned a thickness of 20 μm [61] and a conductivity of 0.06 S/m. Note that the specific conductances of the electrodes do not have a strong influence in our simulations since we directly controlled the current as in the experiments (and the electrodes are not too thick). What is more critical to shape the electric field are the conductance (σ_CSF_) and thickness of the CSF (δ_CSF_), and the conductance of the brain (σ_Brain_). We decided to estimate these parameters based on our experimental data. Specifically, we implemented an optimization procedure to identify the parameters that lead to electric fields that best match our experimental recordings, namely the normal component of the electric field, E_z_, that we recorded at a specific distance from the electrode. We decided to set up the optimization procedure considering not the experimental points directly but the double exponential fit that tracks the spatial profile of the electric field (as in **Figure 4C**). This greatly improves the efficiency of the optimization to converge and allows a direct comparison with the electric field spatial profile that can be fully estimated in the simulations. For the simulations performed within the optimization procedure, we meshed the geometry sufficiently but keeping the meshing not too fine to limit the computational time required for the algorithm to converge. To again facilitate the convergence of the algorithm, we also smoothed the electric field spatial profile in the simulation using a 200 μm window. We defined a cost function as f = Σ_i_ (y_i,sim_ - y_i,data_)^2^ / Σ_i_ y_i,data_^2^ where the index *i* indicates the electric field estimates at different positions for the model (y_sim_) and the experiments fits (y_data_) starting at the first and ending with the last experimentally measured depths (375 μm and 2500 μm). To minimize the cost function we used the default interior-point method of the *fmincon* function in MATLAB where we set the lower and upper bound for the parameters search. The values of the conductances were bounded by the minimum and maximum values reported by Sim4Life (https://itis.swiss/virtual-population/tissue-properties/database/low-frequency-conductivity/), which were derived by a detailed literature review. In **Figure 4G** we also reported the mean and standard deviation of the parameters according to the values reported by Sim4Life. In **Table 1** we report the minimum and maximum values that we considered for the optimization. Note that a previous computational study assumed a thickness of 100 μm for the CSF layer in the mouse [60] but this number likely depends on the specific brain area considered [61]. Thus, we considered the CSF thickness a parameter to optimize together with the conductivities and manually set it to potentially vary in a quite large interval for the optimization procedure. The optimization was performed starting from the mean value of the conductivity parameters as reported in the literature and the average of the min and max values set for the CSF layer thickness. The optimization was repeated 10 times to identify the global minimum of the cost function. Once the optimal values of the parameters were found, we re-run the simulation with the optimal set while further increasing the meshing quality to obtain our final estimation of the electric field distribution around and beneath the stimulation electrode (**Figure 4H,I**).

**Table 1.** Minimum and maximum values of the parameters used for the optimization procedure. Values to optimize are the conductivities of the CSF and the brain as well as the thickness of the CSF layer. Minimum and maximum values for the conductivities are from a review on estimations from the literature by Sim4Life (https://itis.swiss/virtual-population/tissue-properties/database/low-frequency-conductivity/).

| Parameter | Min value | Max value |
| --- | --- | --- |
| $\sigma_{\text{CSF}}$ | 1.13 S/m | 3.19 S/m |
| $\sigma_{\text{Brain}}$ | 0.066 S/m | 0.717 S/m |
| $\delta_{\text{CSF}}$ | 10 $\mu\text{m}$ | 100 $\mu\text{m}$ |

## ACKNOWLEDGEMENTS

We thank Martin Baca for technical support and Fanny Cazettes for helpful comments on the manuscript. This work was funded by Marie-Curie postdoctoral fellowships (D.R.: HORIZON-MSCA-2021-PF-01 101063075), the European Union’s Horizon 2020 research and innovation programme under grant agreement N°101034324 (D.R.), the French government under the France 2030 investment plan, as part of the Initiative d’Excellence d’Aix-Marseille Université – A*MIDEX AMX-22-COF-132 (D.R.), the French National Research Agency, through the « Investissements d’Avenir » program (ANR-21-ESRE-0003).

## AUTHOR CONTRIBUTIONS

Conceptualization: RM, DR, DM, RPO. Methodology: RM, DR. Software: DR. Validation: RM, DR, DM, RPO. Formal Analysis: RM, DR. Investigation: RM, DR, AL. Resources: RM, DR, AL. Data Curation: DR. Writing – Original Draft: DR. Writing – Review & Editing: RM, DR, AL, DM, RPO. Visualization: RM, DR. Supervision: DM, RPO. Project Administration: DR, DM, RPO. Funding Acquisition: DR, DM, RPO.

## SUPPLEMENTARY FIGURES

**Supplementary Figure 1.**
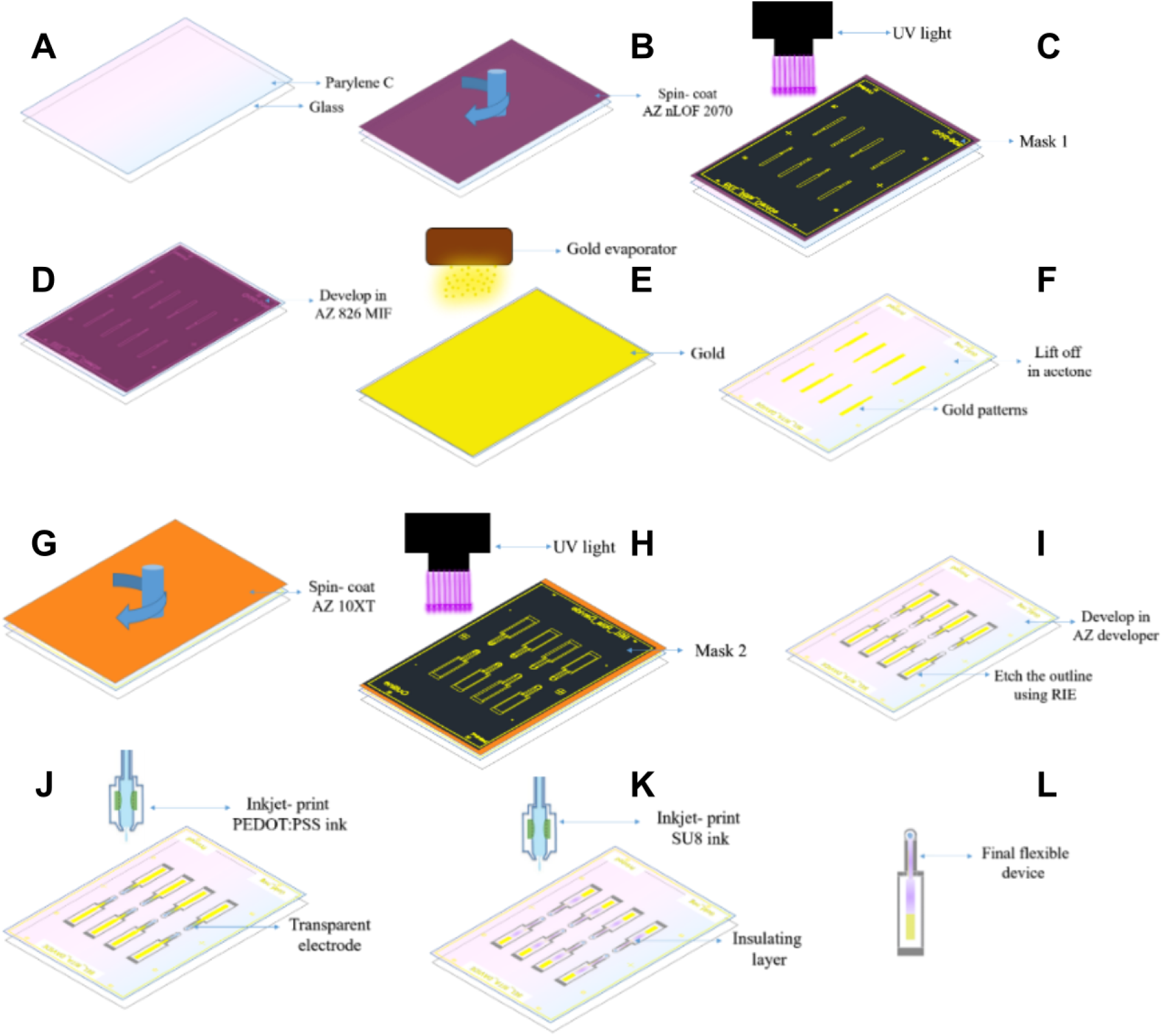
Schematics of the full fabrication process. **A**: PaC deposition. **B**: Spin-coating AZnLOF 2070. **C**: UV Exposure. **D**: Developing. **E**: Gold evaporation. **F**: Lift-off. **G**: Spin-coating AZ 10XT. **H**: UV Exposure. **I**: Developing and reactive ion etching. **J**: Inkjet-printing PEDOT:PSS. **K**: Inkjet-printing SU-8. **L**: Final device.

**Supplementary Figure 2.**
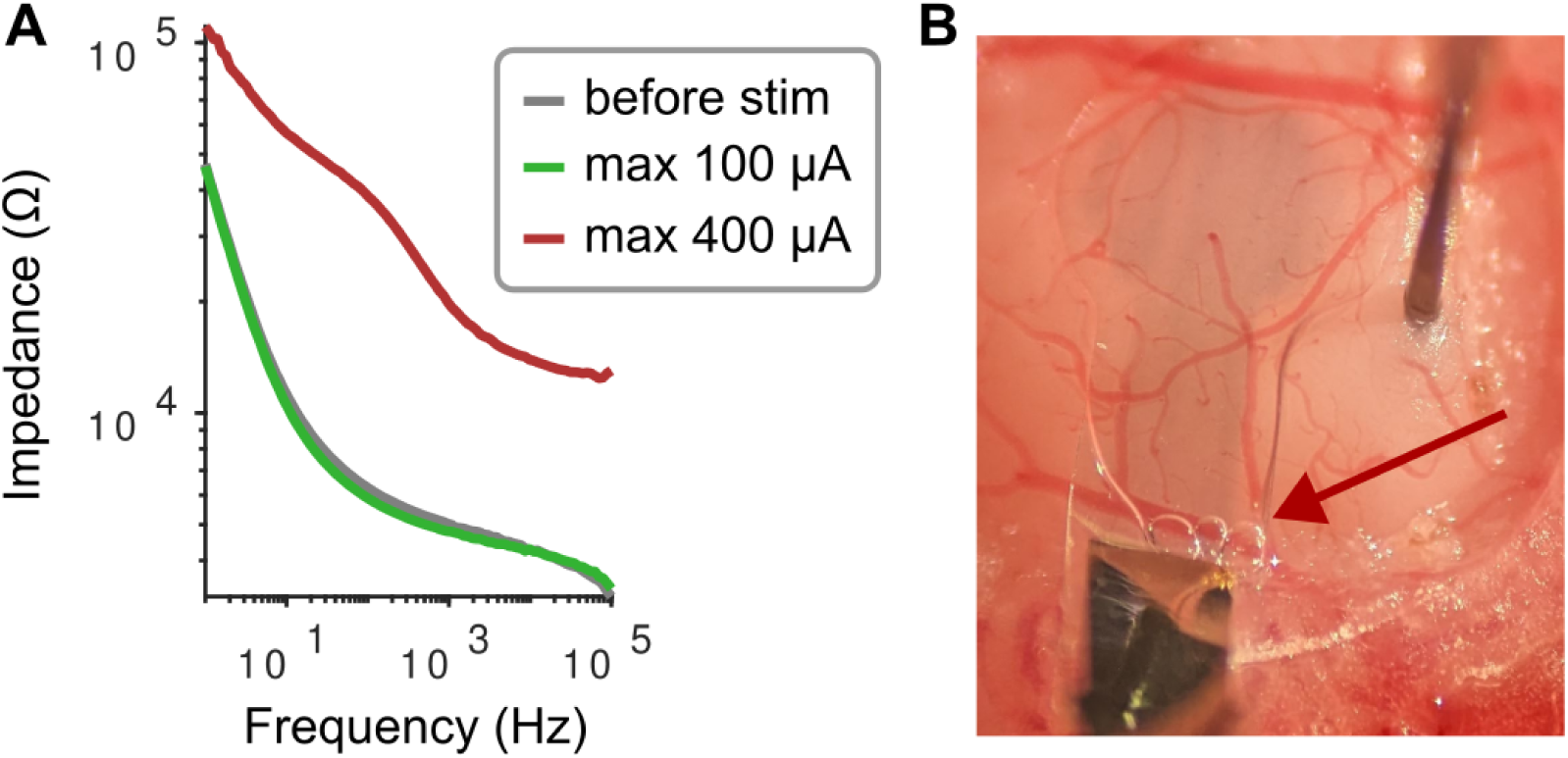
In-vivo testing of electric currents beyond the electrochemically stable water window. **A**: In-vivo electrical impedance spectroscopy measurements before applying the stimulation (gray) after applying a set of stimulation with an amplitude of maximum 100 μA (green) and after a subsequent stimulation set with an amplitude of maximum 400 μA (red). **B**: Occurrence of bubbles (indicated by the red arrow) after the second set of stimulations. Note that they appeared at the intersection between PEDOT:PSS and the gold connector.

## REFERENCES

1. Penfield W, Boldrey E. Somatic motor and sensory representation in the cerebral cortex of man as studied by electrical stimulation. Brain J Neurol. 1937;60: 389–443. doi:10.1093/brain/60.4.389

2. Cohen MR, Newsome WT. What electrical microstimulation has revealed about the neural basis of cognition. Curr Opin Neurobiol. 2004;14: 169–177. doi:10.1016/j.conb.2004.03.016

3. Brunoni AR, Nitsche MA, Bolognini N, Bikson M, Wagner T, Merabet L, et al. Clinical research with transcranial direct current stimulation (tDCS): challenges and future directions. Brain Stimulat. 2012;5: 175–195. doi:10.1016/j.brs.2011.03.002

4. Elyamany O, Leicht G, Herrmann CS, Mulert C. Transcranial alternating current stimulation (tACS): from basic mechanisms towards first applications in psychiatry. Eur Arch Psychiatry Clin Neurosci. 2021;271: 135–156. doi:10.1007/s00406-020-01209-9

5. Perlmutter JS, Mink JW. Deep brain stimulation. Annu Rev Neurosci. 2006;29: 229–257. doi:10.1146/annurev.neuro.29.051605.112824

6. Tehovnik EJ, Slocum WM, Smirnakis SM, Tolias AS. Microstimulation of visual cortex to restore vision. Prog Brain Res. 2009;175: 347–375. doi:10.1016/S0079-6123(09)17524-6

7. Flesher SN, Collinger JL, Foldes ST, Weiss JM, Downey JE, Tyler-Kabara EC, et al. Intracortical microstimulation of human somatosensory cortex. Sci Transl Med. 2016;8: 361ra141. doi:10.1126/scitranslmed.aaf8083

8. Chen X, Wang F, Fernandez E, Roelfsema PR. Shape perception via a high-channel-count neuroprosthesis in monkey visual cortex. Science. 2020;370: 1191–1196. doi:10.1126/science.abd7435

9. McIntyre CC, Mori S, Sherman DL, Thakor NV, Vitek JL. Electric field and stimulating influence generated by deep brain stimulation of the subthalamic nucleus. Clin Neurophysiol Off J Int Fed Clin Neurophysiol. 2004;115: 589–595. doi:10.1016/j.clinph.2003.10.033

10. Huang Y, Liu AA, Lafon B, Friedman D, Dayan M, Wang X, et al. Measurements and models of electric fields in the in vivo human brain during transcranial electric stimulation. eLife. 2017;6. doi:10.7554/eLife.18834

11. Ranck JB. Which elements are excited in electrical stimulation of mammalian central nervous system: a review. Brain Res. 1975;98: 417–440. doi:10.1016/0006-8993(75)90364-9

12. Rattay F. The basic mechanism for the electrical stimulation of the nervous system. Neuroscience. 1999;89: 335–346. doi:10.1016/s0306-4522(98)00330-3

13. Liu A, Vöröslakos M, Kronberg G, Henin S, Krause MR, Huang Y, et al. Immediate neurophysiological effects of transcranial electrical stimulation. Nat Commun. 2018;9: 5092. doi:10.1038/s41467-018-07233-7

14. Purpura DP, Mcmurtry JG. INTRACELLULAR ACTIVITIES AND EVOKED POTENTIAL CHANGES DURING POLARIZATION OF MOTOR CORTEX. J Neurophysiol. 1965;28: 166–185. doi:10.1152/jn.1965.28.1.166

15. Bikson M, Inoue M, Akiyama H, Deans JK, Fox JE, Miyakawa H, et al. Effects of uniform extracellular DC electric fields on excitability in rat hippocampal slices in vitro. J Physiol. 2004;557: 175–190. doi:10.1113/jphysiol.2003.055772

16. Chan CY, Nicholson C. Modulation by applied electric fields of Purkinje and stellate cell activity in the isolated turtle cerebellum. J Physiol. 1986;371: 89–114. doi:10.1113/jphysiol.1986.sp015963

17. Radman T, Ramos RL, Brumberg JC, Bikson M. Role of Cortical Cell Type and Morphology in Sub- and Suprathreshold Uniform Electric Field Stimulation. Brain Stimulat. 2009;2: 215–228. doi:10.1016/j.brs.2009.03.007

18. Rahman A, Reato D, Arlotti M, Gasca F, Datta A, Parra LC, et al. Cellular effects of acute direct current stimulation: somatic and synaptic terminal effects. J Physiol. 2013;591: 2563–2578.

19. Sánchez-León CA, Campos GS-G, Fernández M, Sánchez-López Á, Medina JF, Márquez-Ruiz J. Somatodendritic orientation determines tDCS-induced neuromodulation of Purkinje cell activity in awake mice. bioRxiv; 2023. p. 2023.02.18.529047. doi:10.1101/2023.02.18.529047

20. Terzuolo CA, Bullock TH. MEASUREMENT OF IMPOSED VOLTAGE GRADIENT ADEQUATE TO MODULATE NEURONAL FIRING. Proc Natl Acad Sci U S A. 1956;42: 687–694. doi:10.1073/pnas.42.9.687

21. Radman T, Su Y, An JH, Parra LC, Bikson M. Spike Timing Amplifies the Effect of Electric Fields on Neurons: Implications for Endogenous Field Effects. J Neurosci. 2007;27: 3030–3036. doi:10.1523/JNEUROSCI.0095-07.2007

22. Reato D, Rahman A, Bikson M, Parra LC. Low-intensity electrical stimulation affects network dynamics by modulating population rate and spike timing. J Neurosci. 2010;30: 15067–15079.

23. Farahani F, Khadka N, Parra LC, Bikson M, Vöröslakos M. Transcranial electric stimulation modulates firing rate at clinically relevant intensities. Brain Stimulat. 2024;17: 561–571. doi:10.1016/j.brs.2024.04.007

24. Scala F, Kobak D, Bernabucci M, Bernaerts Y, Cadwell CR, Castro JR, et al. Phenotypic variation of transcriptomic cell types in mouse motor cortex. Nature. 2021;598: 144–150. doi:10.1038/s41586-020-2907-3

25. Peng H, Xie P, Liu L, Kuang X, Wang Y, Qu L, et al. Morphological diversity of single neurons in molecularly defined cell types. Nature. 2021;598: 174–181. doi:10.1038/s41586-021-03941-1

26. Campagnola L, Seeman SC, Chartrand T, Kim L, Hoggarth A, Gamlin C, et al. Local connectivity and synaptic dynamics in mouse and human neocortex. Science. 375: eabj5861. doi:10.1126/science.abj5861

27. Grover S, Nguyen JA, Reinhart RMG. Synchronizing Brain Rhythms to Improve Cognition. Annu Rev Med. 2021;72: 29–43. doi:10.1146/annurev-med-060619-022857

28. Wischnewski M, Alekseichuk I, Opitz A. Neurocognitive, physiological, and biophysical effects of transcranial alternating current stimulation. Trends Cogn Sci. 2023;27: 189–205. doi:10.1016/j.tics.2022.11.013

29. Moro E, Esselink RJA, Xie J, Hommel M, Benabid AL, Pollak P. The impact on Parkinson’s disease of electrical parameter settings in STN stimulation. Neurology. 2002;59: 706–713. doi:10.1212/wnl.59.5.706

30. Opri E, Cernera S, Molina R, Eisinger RS, Cagle JN, Almeida L, et al. Chronic embedded cortico-thalamic closed-loop deep brain stimulation for the treatment of essential tremor. Sci Transl Med. 2020;12: eaay7680. doi:10.1126/scitranslmed.aay7680

31. Scangos KW, Khambhati AN, Daly PM, Makhoul GS, Sugrue LP, Zamanian H, et al. Closed-loop neuromodulation in an individual with treatment-resistant depression. Nat Med. 2021;27: 1696–1700. doi:10.1038/s41591-021-01480-w

32. Bonizzato M, Guay Hottin R, Côté SL, Massai E, Choinière L, Macar U, et al. Autonomous optimization of neuroprosthetic stimulation parameters that drive the motor cortex and spinal cord outputs in rats and monkeys. Cell Rep Med. 2023;4: 101008. doi:10.1016/j.xcrm.2023.101008

33. Zhang Y, Rózsa M, Liang Y, Bushey D, Wei Z, Zheng J, et al. Fast and sensitive GCaMP calcium indicators for imaging neural populations. Nature. 2023;615: 884–891. doi:10.1038/s41586-023-05828-9

34. Liu Z, Lu X, Villette V, Gou Y, Colbert KL, Lai S, et al. Sustained deep-tissue voltage recording using a fast indicator evolved for two-photon microscopy. Cell. 2022;185: 3408–3425.e29. doi:10.1016/j.cell.2022.07.013

35. Lecoq J, Orlova N, Grewe BF. Wide. Fast. Deep: Recent Advances in Multiphoton Microscopy of In Vivo Neuronal Activity. J Neurosci Off J Soc Neurosci. 2019;39: 9042–9052. doi:10.1523/JNEUROSCI.1527-18.2019

36. Madisen L, Garner AR, Shimaoka D, Chuong AS, Klapoetke NC, Li L, et al. Transgenic mice for intersectional targeting of neural sensors and effectors with high specificity and performance. Neuron. 2015;85: 942–958. doi:10.1016/j.neuron.2015.02.022

37. Donahue MJ, Kaszas A, Turi GF, Rózsa B, Slézia A, Vanzetta I, et al. Multimodal Characterization of Neural Networks Using Highly Transparent Electrode Arrays. eNeuro. 2018;5: ENEURO.0187-18.2018. doi:10.1523/ENEURO.0187-18.2018

38. Boehler C, Aqrawe Z, Asplund M. Applications of PEDOT in Bioelectronic Medicine. Bioelectron Med. 2019;2: 89–99. doi:10.2217/bem-2019-0014

39. Cui X, Martin DC. Electrochemical deposition and characterization of poly(3,4-ethylenedioxythiophene) on neural microelectrode arrays. Sens Actuators B Chem. 2003;89: 92–102. doi:10.1016/S0925-4005(02)00448-3

40. Khodagholy D, Gelinas JN, Thesen T, Doyle W, Devinsky O, Malliaras GG, et al. NeuroGrid: recording action potentials from the surface of the brain. Nat Neurosci. 2015;18: 310–315. doi:10.1038/nn.3905

41. Boehler C, Asplund M. PEDOT as a high charge injection material for low-frequency stimulation. 2018 40th Annual International Conference of the IEEE Engineering in Medicine and Biology Society (EMBC). 2018. pp. 2202–2205. doi:10.1109/EMBC.2018.8512597

42. Dijk G, Ruigrok HJ, O’Connor RP. Influence of PEDOT:PSS Coating Thickness on the Performance of Stimulation Electrodes. Adv Mater Interfaces. 2020;7: 2000675. 10.1002/admi.202000675

43. Paulsen BD, Tybrandt K, Stavrinidou E, Rivnay J. Organic mixed ionic-electronic conductors. Nat Mater. 2020;19: 13–26. doi:10.1038/s41563-019-0435-z

44. Orlemann C, Boehler C, Kooijmans RN, Li B, Asplund M, Roelfsema PR. Flexible Polymer Electrodes for Stable Prosthetic Visual Perception in Mice. Adv Healthc Mater. 2024;13: e2304169. doi:10.1002/adhm.202304169

45. Cummins G, Desmulliez MPY. Inkjet printing of conductive materials: a review. Circuit World. 2012;38: 193–213. doi:10.1108/03056121211280413

46. Matta R, Moreau D, O’Connor R. Printable devices for neurotechnology. Front Neurosci. 2024;18. doi:10.3389/fnins.2024.1332827

47. Shi H, Liu C, Jiang Q, Xu J. Effective Approaches to Improve the Electrical Conductivity of PEDOT:PSS: A Review. Adv Electron Mater. 2015;1: 1500017. doi:10.1002/aelm.201500017

48. Galliani M, Ferrari LM, Bouet G, Eglin D, Ismailova E. Tailoring inkjet-printed PEDOT:PSS composition toward green, wearable device fabrication. APL Bioeng. 2023;7: 016101. doi:10.1063/5.0117278

49. Vosgueritchian M, Lipomi DJ, Bao Z. Highly Conductive and Transparent PEDOT:PSS Films with a Fluorosurfactant for Stretchable and Flexible Transparent Electrodes. Adv Funct Mater. 2012;22: 421–428. doi:10.1002/adfm.201101775

50. Dijk G, Kaszas A, Pas J, O’Connor RP. Fabrication and in vivo 2-photon microscopy validation of transparent PEDOT:PSS microelectrode arrays. Microsyst Nanoeng. 2022;8: 1–8. doi:10.1038/s41378-022-00434-7

51. Fekete Z, Zátonyi A, Kaszás A, Madarász M, Slézia A. Transparent neural interfaces: challenges and solutions of microengineered multimodal implants designed to measure intact neuronal populations using high-resolution electrophysiology and microscopy simultaneously. Microsyst Nanoeng. 2023;9: 1–30. doi:10.1038/s41378-023-00519-x

52. Nguyen VH, Papanastasiou DT, Resende J, Bardet L, Sannicolo T, Jiménez C, et al. Advances in Flexible Metallic Transparent Electrodes. Small. 2022;18: 2106006. doi:10.1002/smll.202106006

53. Bae S, Kim H, Lee Y, Xu X, Park J-S, Zheng Y, et al. Roll-to-roll production of 30-inch graphene films for transparent electrodes. Nat Nanotechnol. 2010;5: 574–578. doi:10.1038/nnano.2010.132

54. Kuzum D, Takano H, Shim E, Reed JC, Juul H, Richardson AG, et al. Transparent and flexible low noise graphene electrodes for simultaneous electrophysiology and neuroimaging. Nat Commun. 2014;5: 5259. doi:10.1038/ncomms6259

55. Park D-W, Ness JP, Brodnick SK, Esquibel C, Novello J, Atry F, et al. Electrical Neural Stimulation and Simultaneous in Vivo Monitoring with Transparent Graphene Electrode Arrays Implanted in GCaMP6f Mice. ACS Nano. 2018;12: 148–157. doi:10.1021/acsnano.7b04321

56. Torrisi F, Hasan T, Wu W, Sun Z, Lombardo A, Kulmala TS, et al. Inkjet-Printed Graphene Electronics. ACS Nano. 2012;6: 2992–3006. doi:10.1021/nn2044609

57. Li J, Ye F, Vaziri S, Muhammed M, Lemme MC, Östling M. Efficient Inkjet Printing of Graphene. Adv Mater. 2013;25: 3985–3992. doi:10.1002/adma.201300361

58. McIntyre CC, Grill WM. Finite Element Analysis of the Current-Density and Electric Field Generated by Metal Microelectrodes. Ann Biomed Eng. 2001;29: 227–235. doi:10.1114/1.1352640

59. Datta A, Bansal V, Diaz J, Patel J, Reato D, Bikson M. Gyri-precise head model of transcranial direct current stimulation: improved spatial focality using a ring electrode versus conventional rectangular pad. Brain Stimulat. 2009;2: 201–207.

60. Alekseichuk I, Mantell K, Shirinpour S, Opitz A. Comparative modeling of transcranial magnetic and electric stimulation in mouse, monkey, and human. NeuroImage. 2019;194: 136–148. doi:10.1016/j.neuroimage.2019.03.044

61. Møllgård K, Beinlich FRM, Kusk P, Miyakoshi LM, Delle C, Plá V, et al. A mesothelium divides the subarachnoid space into functional compartments. Science. 2023;379: 84–88. doi:10.1126/science.adc8810

62. Deng Z-D, Lisanby SH, Peterchev AV. Electric field strength and focality in electroconvulsive therapy and magnetic seizure therapy: a finite element simulation study. J Neural Eng. 2011;8: 016007. doi:10.1088/1741-2560/8/1/016007

63. Luo Y, Abidian MR, Ahn J-H, Akinwande D, Andrews AM, Antonietti M, et al. Technology Roadmap for Flexible Sensors. ACS Nano. 2023;17: 5211–5295. doi:10.1021/acsnano.2c12606

64. Qiu J, Yu X, Wu X, Wu Z, Song Y, Zheng Q, et al. An Efficiently Doped PEDOT:PSS Ink Formulation via Metastable Liquid−Liquid Contact for Capillary Flow-Driven, Hierarchically and Highly Conductive Films. Small. 2023;19: 2205324. doi:10.1002/smll.202205324

65. Ahmad Shahrim N, Ahmad Z, Azman AW, Buys YF, Sarifuddin N. Mechanisms for doped PEDOT:PSS electrical conductivity improvement. Mater Adv. 2021;2: 7118–7138. doi:10.1039/D1MA00290B

66. Kim S, Kim SY, Kim J, Kim JH. Highly reliable AgNW/PEDOT:PSS hybrid films: efficient methods for enhancing transparency and lowering resistance and haziness. J Mater Chem C. 2014;2: 5636–5643. doi:10.1039/C4TC00686K

67. Ganji M, Hossain L, Tanaka A, Thunemann M, Halgren E, Gilja V, et al. Monolithic and Scalable Au Nanorod Substrates Improve PEDOT-Metal Adhesion and Stability in Neural Electrodes. Adv Healthc Mater. 2018;7: e1800923. doi:10.1002/adhm.201800923

68. Smołka S, Skorupa M, Barylski A, Basiaga M, Krukiewicz K. Improved adhesion and charge transfer between PEDOT:PSS and the surface of a platinum electrode through a diazonium chemistry route. Electrochem Commun. 2023;153: 107528. doi:10.1016/j.elecom.2023.107528

69. McIntyre CC, Grill WM. Current density and electric field analysis of microelectrodes using finite element modeling. Proceedings of the 22nd Annual International Conference of the IEEE Engineering in Medicine and Biology Society (Cat No00CH37143). 2000. pp. 2010–2011 vol.3. doi:10.1109/IEMBS.2000.900491

70. Weaver JC, Smith KC, Esser AT, Son RS, Gowrishankar TR. A brief overview of electroporation pulse strength-duration space: A region where additional intracellular effects are expected. Bioelectrochemistry Amst Neth. 2012;87: 236–243. doi:10.1016/j.bioelechem.2012.02.007

71. Guillen A, Abbott CC, Deng Z-D, Huang Y, Pascoal-Faria P, Truong DQ, et al. Impact of modeled field of view in electroconvulsive therapy current flow simulations. Front Psychiatry. 2023;14: 1168672. doi:10.3389/fpsyt.2023.1168672

72. Rampersad S, Roig-Solvas B, Yarossi M, Kulkarni PP, Santarnecchi E, Dorval AD, et al. Prospects for transcranial temporal interference stimulation in humans: A computational study. NeuroImage. 2019;202: 116124. doi:10.1016/j.neuroimage.2019.116124

73. Liu R, Zhu G, Wu Z, Gan Y, Zhang J, Liu J, et al. Temporal interference stimulation targets deep primate brain. NeuroImage. 2024;291: 120581. doi:10.1016/j.neuroimage.2024.120581

74. Rudiak D, Marg E. Finding the depth of magnetic brain stimulation: a re-evaluation. Electroencephalogr Clin Neurophysiol Potentials Sect. 1994;93: 358–371. doi:10.1016/0168-5597(94)90124-4

75. Thielscher A, Kammer T. Linking Physics with Physiology in TMS: A Sphere Field Model to Determine the Cortical Stimulation Site in TMS. NeuroImage. 2002;17: 1117–1130. doi:10.1006/nimg.2002.1282

76. Reato D, Salvador R, Bikson M, Opitz A, Dmochowski J, Miranda PC. Principles of transcranial direct current stimulation (tDCS): introduction to the biophysics of tDCS. Practical guide to transcranial direct current stimulation. Springer; 2019. pp. 45–80.

77. Ozen S, Sirota A, Belluscio MA, Anastassiou CA, Stark E, Koch C, et al. Transcranial electric stimulation entrains cortical neuronal populations in rats. J Neurosci Off J Soc Neurosci. 2010;30: 11476–11485. doi:10.1523/JNEUROSCI.5252-09.2010

78. Anastassiou CA, Perin R, Markram H, Koch C. Ephaptic coupling of cortical neurons. Nat Neurosci. 2011;14: 217–223. doi:10.1038/nn.2727

79. Krause MR, Vieira PG, Csorba BA, Pilly PK, Pack CC. Transcranial alternating current stimulation entrains single-neuron activity in the primate brain. Proc Natl Acad Sci U S A. 2019;116: 5747–5755. doi:10.1073/pnas.1815958116

80. Fröhlich F, McCormick DA. Endogenous Electric Fields May Guide Neocortical Network Activity. Neuron. 2010;67: 129–143. doi:10.1016/j.neuron.2010.06.005

81. Johnson L, Alekseichuk I, Krieg J, Doyle A, Yu Y, Vitek J, et al. Dose-dependent effects of transcranial alternating current stimulation on spike timing in awake nonhuman primates. Sci Adv. 2020;6. doi:10.1126/sciadv.aaz2747

82. Creutzfeldt OD, Fromm GH, Kapp H. Influence of transcortical d-c currents on cortical neuronal activity. Exp Neurol. 1962;5: 436–452. doi:10.1016/0014-4886(62)90056-0

83. Bindman LJ, Lippold OC, Redfearn JW. THE ACTION OF BRIEF POLARIZING CURRENTS ON THE CEREBRAL CORTEX OF THE RAT (1) DURING CURRENT FLOW AND (2) IN THE PRODUCTION OF LONG-LASTING AFTER-EFFECTS. J Physiol. 1964;172: 369–382. doi:10.1113/jphysiol.1964.sp007425

84. Histed MH, Bonin V, Reid RC. Direct activation of sparse, distributed populations of cortical neurons by electrical microstimulation. Neuron. 2009;63: 508–522. doi:10.1016/j.neuron.2009.07.016

85. Trevathan JK, Asp AJ, Nicolai EN, Trevathan JM, Kremer NA, Kozai TD, et al. Calcium imaging in freely moving mice during electrical stimulation of deep brain structures. J Neural Eng. 2020; 10.1088/1741-2552/abb7a4. doi:10.1088/1741-2552/abb7a4

86. Eles JR, Stieger KC, Kozai TDY. The temporal pattern of intracortical microstimulation pulses elicits distinct temporal and spatial recruitment of cortical neuropil and neurons. J Neural Eng. 2021;18: 015001. doi:10.1088/1741-2552/abc29c

87. Wu GK, Ardeshirpour Y, Mastracchio C, Kent J, Caiola M, Ye M. Amplitude- and frequency-dependent activation of layer II/III neurons by intracortical microstimulation. iScience. 2023;26. doi:10.1016/j.isci.2023.108140

88. Dadarlat MC, Sun YJ, Stryker MP. Activity-dependent recruitment of inhibition and excitation in the awake mammalian cortex during electrical stimulation. Neuron. 2024;112: 821–834.e4. doi:10.1016/j.neuron.2023.11.022

89. Vöröslakos M, Takeuchi Y, Brinyiczki K, Zombori T, Oliva A, Fernández-Ruiz A, et al. Direct effects of transcranial electric stimulation on brain circuits in rats and humans. Nat Commun. 2018;9: 483. doi:10.1038/s41467-018-02928-3

90. Lee SY, Kozalakis K, Baftizadeh F, Campagnola L, Jarsky T, Koch C, et al. Cell-class-specific electric field entrainment of neural activity. Neuron. 2024;0. doi:10.1016/j.neuron.2024.05.009

91. Chakraborty D, Truong DQ, Bikson M, Kaphzan H. Neuromodulation of Axon Terminals. Cereb Cortex N Y N 1991. 2018;28: 2786–2794. doi:10.1093/cercor/bhx158

92. Bando Y, Wenzel M, Yuste R. Simultaneous two-photon imaging of action potentials and subthreshold inputs in vivo. Nat Commun. 2021;12: 7229. doi:10.1038/s41467-021-27444-9

93. Kannan M, Vasan G, Haziza S, Huang C, Chrapkiewicz R, Luo J, et al. Dual-polarity voltage imaging of the concurrent dynamics of multiple neuron types. Science. 2022;378: eabm8797. doi:10.1126/science.abm8797

94. Joucla S, Yvert B. The “mirror” estimate: an intuitive predictor of membrane polarization during extracellular stimulation. Biophys J. 2009;96: 3495–3508. doi:10.1016/j.bpj.2008.12.3961

95. Aberra AS, Peterchev AV, Grill WM. Biophysically realistic neuron models for simulation of cortical stimulation. J Neural Eng. 2018;15: 066023. doi:10.1088/1741-2552/aadbb1

96. Tran H, Shirinpour S, Opitz A. Effects of transcranial alternating current stimulation on spiking activity in computational models of single neocortical neurons. NeuroImage. 2022; 118953. doi:10.1016/j.neuroimage.2022.118953

